# Reduced offspring viability is associated with long-term stability of a narrow avian hybrid zone

**DOI:** 10.1101/2025.05.22.655562

**Authors:** Kira M. Long, Michael J. Braun, Adolfo Muñoz Abrego, Ovidio Jaramillo, Jeffrey D. Brawn

**Author notes:** Corresponding authors contact details: Kira M. Long and Jeffrey D. Brawn.

## Abstract

Fitness of hybrid individuals can shape the dynamics of hybrid zones and offer insight into speciation processes. Yet, accounts of hybrid fitness in natural hybrid zones are few, especially from tropical regions where species diversity is high, and speciation processes could contrast with those at higher latitudes. We investigated a hybrid zone between the white-collared manakin (*Manacus candei*) and the golden-collared manakin (*M. vitellinus*), two lek-breeding species characteristic of lowland forest habitat in Central America. Despite evidence of asymmetrical introgression and selection on male secondary sexual traits, ongoing sampling indicates that this hybrid zone is spatially stable with narrow clines, thus implying selection against hybrids. To evaluate hybrid viability, we estimated two components of hybrid fitness: survival of adults and egg hatching rates, and a possible selective pressure: prevalence of parasitism by vector-borne haemosporidian parasites. Estimated survival was similar between parental and hybrid populations and the prevalence of infections by *Plasmodium* spp. or *Haemoproteus* spp. parasites was uniformly low. Estimated rates of hatching success, however, were lower in nests from our hybrid population (one or two eggs failed to hatch in 70% of nests) compared to nests at the parental species (*M. candei* 28.6% and *M. vitellinus* 19.0%). Thus, despite extensive admixture and clear evidence of introgression of male plumage traits under sexual selection, partial infertility or elevated rates of developmental mortality in hybrid offspring may underlie long-term stability in this hybrid zone.

## Introduction

Hybridization is a key evolutionary event that invites inquiry into the processes that generate and maintain species diversity. Hybrid zones form at the overlapping edges of two or more species’ ranges and are dynamic systems that can move across a landscape or remain spatially stable. If hybrids have a selective advantage, they are expected to expand into the ranges of one or both parental species (Buggs 2007). Alternatively, hybrid zones can remain narrow and spatially stable if hybrids have low fitness compared to the parental species (Barton and Hewitt 1985). Thus, hybrid fitness can be key to understanding hybridization dynamics and the processes underlying speciation. Yet, accounts of hybrid fitness in natural hybrid zones are few, especially from tropical regions where species diversity is high and speciation rates and processes could contrast with those at higher latitudes (Weir and Schluter 2007, Weir and Price 2011).

Reduced viability of hybrids and consequent fitness costs to parental species can be realized through a variety of mechanisms across taxa, including reduced survival rates and fertility in fish (Gilk et al. 2004, Muhlfeld et al. 2009, Stelkens et al. 2015) and mammals (Lancaster et al. 2007, White et al. 2012, Adavoudi and Pilot 2021). For birds, higher rates of hatching failure have been reported in both outbred and inbred populations (Sætre et al. 1999, Bronson et al. 2005, Neubauer et al. 2014, Maxwell et al. 2021, Ålund et al. 2023), suggesting that hatching success could be an important indicator of underlying genetic incompatibilities. Furthermore, mating systems that result in highly skewed reproductive success, such as lek-breeding systems, invite exploration of the tension between sexual selection and natural selection in hybrid zones and possible effects on hybrid fitness.

We assessed the viability of hybrids from two species of lek-breeding birds, the golden-collared manakin (*Manacus vitellinus*) and the white-collared manakin (*Manacus candei*), which hybridize in western Panama. Male manakins perform elaborate mating displays for females seeking mates (Kirwan and Green 2011, Day et al. 2021), with most copulations going to few males (Lill 1974, Emlen and Oring 1977, Payne 1984). In this hybrid zone, female *Manacus* prefer males with yellow collar plumage (Stein and Uy 2006), resulting in the observed introgression of yellow *M. vitellinus* plumage traits for more than 50 km beyond the genomic transition from *vitellinus* to *candei* (Parsons et al. 1993, Brumfield et al. 2001) (Figure 1). Despite multiple lines of evidence that sexual selection favors males with yellow collars (McDonald et al. 2001, Stein and Uy 2006), the genomic transition between the two parental species has remained spatially stable and narrow for at least 30 years, while the darker olive belly color of *M. vitellinus* has continued to introgress into populations that are genomically like *M. candei*, indicating a decoupling of sexual and natural selection (Long et al. 2024).

**Figure 1.**
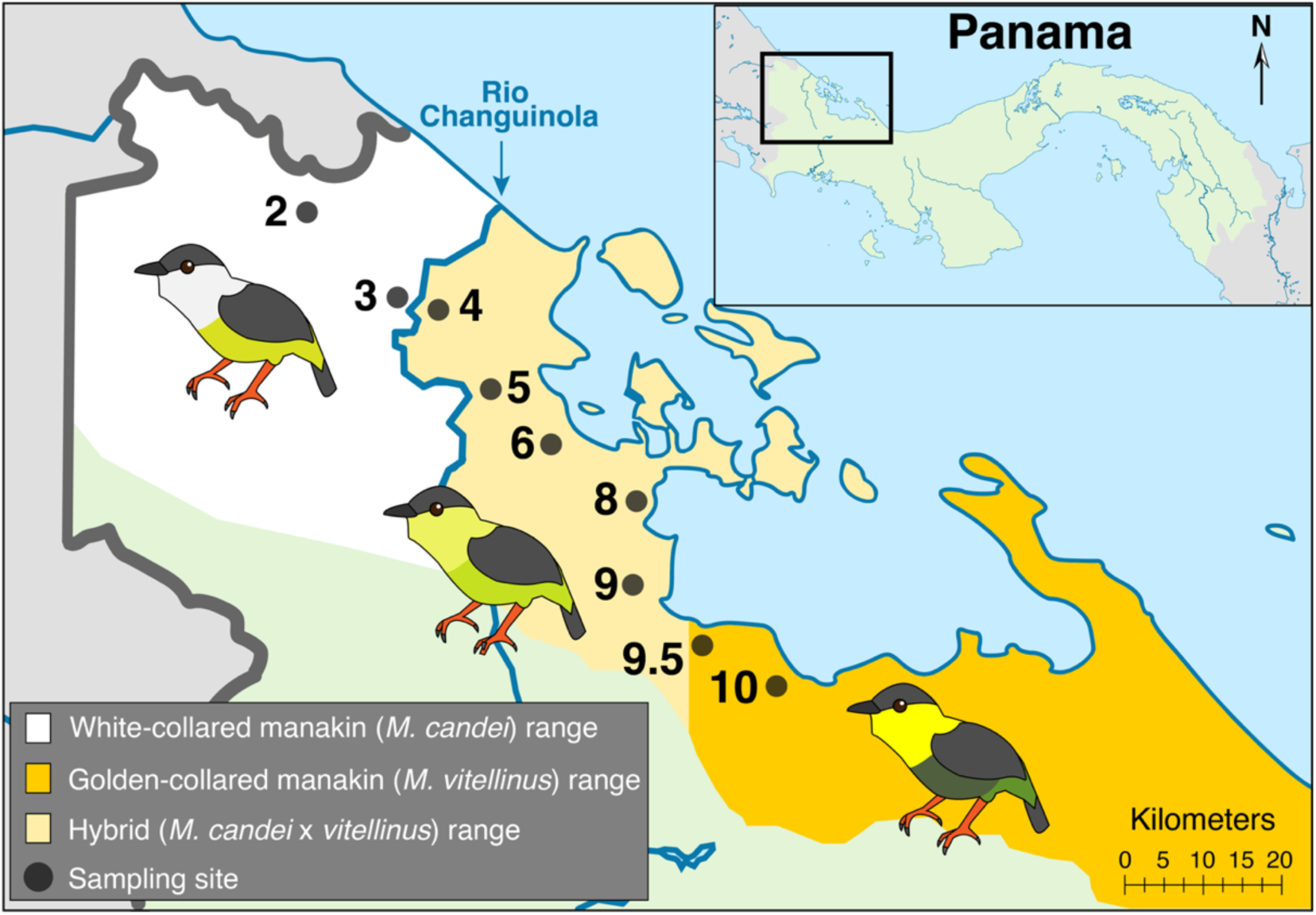
Study region of the *Manacus* hybrid zone in Bocas del Toro, Panama. The black dots represent the sampling sites and are numbered according to Brumfield et al. (2001) and Long et al. (2024). Sampling site 2 is parental *M. candei*, site 10 is parental *M. vitellinus*, and site 9 is the genomic center of the hybrid zone with the most highly admixed hybrid individuals. Note that the hybrid center is at site 9, while the phenotypic transition where yellow collar plumage switches to white collar plumage occurs at the Río Changuinola between sites 3 and 4. Figure 1 has been reproduced from Long et al. (2024).

The spatial stability of this hybrid zone within a narrow genomic transition despite strong sexual selection and ongoing introgression of male secondary sexual traits, suggests the influence of selective pressures against hybrids; specifically, reduced fitness owing to lower adult survival, reduced reproductive output, or both. The specific selective pressure(s) causing reduced hybrid fitness could be direct effects leading to hybrid sterility (White et al. 2012) and offspring inviability (Bolnick and Near 2005), or more subtle factors such as susceptibility to disease and parasites (Goldberg et al. 2005), or physiological impairments like increased oxidative stress (Barreto and Burton 2013, Du et al. 2017, Petrović et al. 2023), which, for male manakins, could particularly compromise their performance in a highly competitive lek-breeding system (Chen et al. 2024).

Parasitism from vector-borne blood parasites can be a strong selective force in wild bird populations (Lachish et al. 2011), especially in immunologically naïve populations when novel parasites are introduced like has been observed with avian malaria (LaPointe et al. 2012, Fecchio et al. 2019, Atkinson 2023). Compromised immunocompetence may be especially disadvantageous for hybrids if they inherit susceptibility to parasites from both parental species (Derothe et al. 2001, Krasnovyd et al. 2017). For manakins, there is evidence of reduced fitness from avian malarial infections causing impaired mating behavior on the lek such as decreased vocalization and display rates (Bosholn et al. 2016). Furthermore, the polygynous lek-breeding system in manakins may increase exposure to parasites and their vectors when individuals gather for breeding displays (Tella 2002, Fecchio et al. 2011). Thus, given strong selection on male traits, the structure of lek-breeding systems, and the expectation that a significant parasite load will negatively impact male secondary sexual traits (Hamilton and Zuk 1982), parasites could be an important selective pressure compromising hybrid fitness in the *Manacus* hybrid system.

To assess whether hybrids have reduced fitness in the *Manacus* hybrid zone, we estimated survival rates of adults and hatching rates of eggs as proxies of hybrid fitness. Given the apparent genomic stability of the *Manacus* hybrid zone, we predicted that adults in the genomic center of the hybrid zone would experience either reduced survival rates and/or nests at the genomic transition would have reduced hatching success when compared to the parental species populations. We also estimated the prevalence of haemosporidian infection (i.e., avian malarial and malarial-like vector-borne blood parasites *Plasmodium* spp. and *Haemoproteus* spp.) as a possible selective pressure on *Manacus* hybrids. We predicted that hybrid *Manacus* individuals would have greater prevalence of blood parasites than the parental species.

## Methods

### Study species and sampling sites

*Manacus vitellinus* and *Manacus candei* (Pipridae) are frugivorous, neotropical birds characteristic of secondary or edge habitats in lowland humid forests. *M. vitellinus* occurs in Panama and western Colombia, while *M. candei* occurs from the Yucatan to Panama. These species’ ranges overlap in Bocas del Toro, Panama, where they are known to hybridize. *Manacus* males congregate at leks where they perform elaborate courtship displays for visiting females (Lill 1974, Kirwan and Green 2011, Day et al. 2021). After mating, only females provide parental care. Both species and their hybrids build open cup nests.

We sampled at 9 sites in a transect across Bocas del Toro, Panama (Figure 1) that were assessed previously by Brumfield et al. (2001) and by Long et al. (2024) (Figure 1). Sites 2 and 10 represent the focal parental species sites for parental *M. candei* and *M. vitellinus*, respectively. Sites 3 and 9.5 represent additional parental sites and all other sites correspond to populations with various levels of parental admixture that we refer to as the “hybrid populations”. Site 9, in particular, represents the hybrid center and is the genomic transition between the two parental populations, displaying the largest variation in parental ancestry, hereafter referred to as the hybrid focal population (Brumfield et al. 2001, Long et al. 2024). Logistics restricted us tosample at focal sites 2, 9, and 10 for estimating annual survival and hatching success, while all 9 sampling sites were assessed for prevalence of haemosporidian infections.

### Capture, mark, and release of individuals

Birds were captured with mist nets in 2017 – 2020 from late February to June. We set up mist nets (36-mm mesh “ATX” type) on leks every 3 days near the displaying males’ courtship arenas to capture displaying males, floating juveniles, and visiting females. We mist netted at one focal site each day, rotating between the three sites so that each site was sampled twice a week.

Nets were checked every 30 minutes from 0800 to 1700 hr and all captured manakins were given an aluminum band with a unique identification number and a combination of plastic colored leg bands for individual identification after release (Raw capture record: Table S1). All protocols involving the capture and handling of live birds were reviewed and approved by the Illinois Animal Care and Use Committee and the Smithsonian Tropical Research Institute Animal Care and Use Committee (Illinois IACUC numbers: 15234 & 18239; STRI ACUC numbers: 2016-0301-2019 & 2019-0115-2022).

### Sex Determination

Sex and reproductive status were determined in the field using definitive adult male plumage or the presence of a brood patch (only females incubate). However, immature males and non-breeding females look nearly identical, with overall olive plumage and yellow bellies, thus their sex was confirmed at a later time using an avian polymerase chain reaction (PCR) molecular sexing protocol that targets an intron of the *CHD1* gene on the Z and W chromosomes (Fridolfsson and Ellegren 1999). Briefly, we extracted blood samples of <100 µL / bird by puncturing the brachial vein. Blood was stored in Longmire’s lysis buffer (Longmire et al. 1997) and kept frozen at -20°C until laboratory analysis. We used primers 2550F and 2718R (2550F = 5’-GTT ACT GAT TCG TCT ACG AGA-3’; 2718R = 5’-ATT GAA ATG ATC CAG TGC TTG-3’) in a master mix with TaKaRa Ex Taq DNA Polymerase reagents (Takara Bio USA, Inc) to perform the PCR outlined in Table A1. The PCR products were visualized for scoring after electrophoresis on a 2% agarose gel stained with GelRed Nucleic Acid Gel Stain (Biotium, Fremont, CA, USA). The targeted Z chromosome region was a 600 bp fragment that would be present in both sexes, while only females had a second 450 bp amplified fragment from the W chromosome (Molecular sexing results: Table S2).

### Estimating adult survival

We rotated mistnetting at the three focal sites each day so that each site was sampled twice a week for the entire five-month field season, resulting in similar sampling effort at each focal site for each sampling year. In 2020, however, we were unable to continue mistnetting operations for the entire 5-month field season due to COVID-19, and thus all mark-recapture data from 2020 were dropped from subsequent analyses because sampling effort was too low. We calculated apparent survival using Cormack-Jolly-Seber (CJS) models for open populations using *Program MARK* (White and Burnham 1999). CJS models calculate apparent survival as the product of the probability of true survival and fidelity to the study area (Lebreton et al. 1992).

Since mortality and emigration cannot be distinguished, apparent survival is an underestimate of true survival (Schaub and Royle 2014). For each individual, we created an encounter history using a custom python script (MARK_extract_input_file). Briefly, this script takes a list of each date a bird was captured, the sampling site it was captured at, and its unique band ID, and creates a year-by-year capture record for that individual and exports an appropriately formatted input file (.inp) for *Program MARK*. For example, if a bird was captured in a 2017, not recaptured in 2018, but recaught in 2019, it would be denoted as ‘101’ for the 2017, 2018, and 2019 field seasons, respectively.

*Program MARK* then provides estimates of apparent survival (denoted as *φ*) and accounts for possible heterogeneity in recapture probability (i.e., probability of recapturing an individual given presence in the sampling area, denoted as *p*) among groups. We compared estimates of apparent survival among white-collared manakins (parental species sampling site 2), golden-collared manakins (parental species sampling site 10), and hybrid individuals at the genomic center of the hybrid zone (hybrid sampling site 9). To explore possible differences in apparent survival probability (*φ*) based on hybrid status (*h*), we used hybrid/parental assignment as a covariate to estimate among the three groups (*φ(h)*). We also accounted for heterogeneity among the three groups in recapture probability (*p*) thus we used the following model in *MARK* to estimate adult survival rate: (*φ(h)p(h)*). Finally, to determine if there were differences in survival between the three groups, we used the apparent survival point estimates and evaluated if their 95% confidence intervals overlapped.

### Hatching Success

We systematically searched for nests in the understory at leks and surrounding forest every 3 days. We also used radio telemetry to locate nests by fitting females with transmitters (PicoPip Ag376 Tags from Lotek Wireless Inc, Canada). The transmitters weighed less than 5% of the birds’ body weight (as recommended by Brander and Cochran 1969, Naef-Daenzer et al. 2001, Barron et al. 2010), and were attached using the Rappole harness technique (Rappole and Tipton 1991). At each located active nest, we “candled” eggs with a flashlight to inspect the interior of the egg every three days to track egg development (Birkhead et al. 2008). Briefly, eggs candled right after being laid will glow yellow when illuminated and can have an observably distinct yolk. If an egg has been fertilized, it will further develop red vasculature, and the egg will glow red when illuminated. As the egg is incubated a developing embryo can be observed, usually appearing as a small, dark shadow that grows until completely eclipsing the interior of the egg, except for the air cell which also grows larger and becomes increasing slanted as incubation continues up until hatching. If a female was found currently sitting on the nest at a three-day nest check, we did not flush the female and skipped candling the eggs that day to reduce the likelihood of nest abandonment. Given that fertilized eggs can fail within the first few hours after fertilization and would appear nearly identical in phenotype to truly unfertilized eggs when using a field candling method, we focus our analyses on eggs that hatched or failed to hatch by the end of the incubation period. To accurately determine if an egg failed due to fertilization failure versus early embryo mortality would require microscopy of egg contents (Assersohn et al. 2021). Nests that failed due to extrinsic factors such as predation, flooding, or human activity such as logging or herding cattle were not included in our analyses. Thus, we defined an egg as failing to hatch based on the definition of hatching failure provided by Marshall et al. 2023; i.e., the nest was not abandoned or destroyed by extrinsic factors such as predation, accident, or extreme weather, and the egg failed to hatch by the end of the expected incubation period (18 days for *M. vitellinus* (Brawn et al. 2011), and we estimated a similar incubation period for *M. candei* or hybrids). To our knowledge, all eggs in our analyses were incubated (Hatching data: Table S3).

To compare hatching rates between the focal parental and hybrid sampling sites, we performed two statistical tests. First, we calculated the proportion of eggs that failed to hatch relative to all eggs that were laid in each focal group and then performed a two-tailed Fisher’s Exact Test for count data using the Freeman-Halton extension to test a 2 × 3 contingency table to compare the three groups using fisher.test() in base R (R Core Team 2022). Second, to account for the non-independence of eggs within the same nest, we fit the hatching success data to a binomial regression, using the success/failure to hatch for each egg within a single nest to a generalized linear model with a binomial distribution. We used the glm() function with the binomial family available in base R, and structured our data so that each row is a nest, with response columns for the success/failure of each egg in the nest (R Core Team 2022). This method counts a binomial ‘success or failure’ outcome for one or more eggs within a nest and thus avoids non-independence of eggs within a single nest.

### Haemosporidian Assays

We tested for avian blood parasites in 268 individuals (Malaria data: Table S4). DNA was extracted from whole blood using an AutoGenprep 965 and the quantity of DNA was determined using a Qubit BR dsDNA assay (Life Technologies), as described in Long et al. (2024). Each sample was screened at least twice to reduce false positives in the dataset. We screened each blood sample for the presence or absence of the vector-borne avian malaria parasites and malaria-like parasites from the genera *Plasmodium* or *Haemoproteus*, respectively; hereafter, haemosporidia is used to refer to both assayed genera. We used PCR to target a highly conserved region of the parasites’ mitochondrial DNA, the 154-bp *16S rRNA* coding sequence. Note that this PCR test will result in a positive result for the presence of either parasitic lineage, and does not differentiate if the infection is *Plasmodium* spp. or *Haemoproteus* spp. We used the primers *343F* and *496R* from Fallon et al. (2003) with the following sequences: *343F* = 5’-GCT CAC GCA TCG CTT CT -3’ and *496R* = 5’-GAC CGG TCA TTT TCT TTG -3’ to perform the PCR outlined in Table A2. The PCR products were visualized for scoring after electrophoresis on a 2% agarose gel stained with GelRed Nucleic Acid Gel Stain (Biotium, Fremont, CA, USA).

Samples were always run with two positive controls to corroborate positive results, and two negative controls to detect any possible contamination, but no contamination was found in any PCR run. All samples were then screened a second time with the same protocol to reduce false positive results. Samples that were positive in the screening PCR were then advanced to a second nested PCR assay that targets the *cytochrome b* gene in the parasites’ mitochondria. For this nested PCR assay, we used the *HAEM* primers from Waldenstrom et al. (2002): *HAEMNF* = 5’ -CAT ATA TTA AGA GAA TTA TGG AG -3’, *HAEMNR2* = 5’ -AGA GGT GTA GCA TAT CTA TCT AC -3’, *HAEMF* = 5’ - ATG GTG CTT TCG ATA TAT GCA TG -3’, and *HAEMR2* = 5’ - GCA TTA TCT GGA TGT GAT AAT GGT -3’ to perform the PCR outlined in Table A3. Samples that were positive in all three PCR assays (twice in the screening step and once in the Waldenstrom step, for three PCR tests total) were considered positive in the final dataset.

## Results

### Hatching Success

We located 47 active nests across the three focal sampling sites. Nine nesting attempts failed (19%) prior to hatching (6 (*M. candei*), 3 (hybrid), and 0 (*M. vitellinus*)). Of the remaining 38 nests, we monitored the developmental progress of eggs throughout the incubation period in 21 (*M. candei*), 10 (hybrid), and 7 (*M. vitellinus)* nests. All nests contained 1-2 eggs at all three focal sites. Six of the 38 nests (16%) contained only one egg for the entire incubation period (4, 1, and 1 *M. candei*, hybrids, and *M. vitellinus,* respectively). Of all the nests with singleton eggs, 2 (33%) failed to hatch. One or both eggs failed to hatch in 11 of 32 (34%) of nests with two eggs. Overall, the proportion of failed eggs from the total eggs laid within each focal site was 13% (*M. candei*, 5 of 38), 42% (hybrid, 8 of 19), and 23% (*M. vitellinus*, 3 of 13) (Fisher’s Exact Test, p-value = 0.0473). Grouping hatching failure by nest to account for the non-independence of eggs, 70% of nests from the hybrid focal population had one or both eggs fail to hatch, a proportion significantly greater than *M. vitellinus* (28.6%) or *M. candei* (19.0%) (Figure 2.; GLM Binomial, Wald statistic z value = -2.348, df = 37, S.E. = 0.67, p-value = 0.0189).

**Figure 2.**
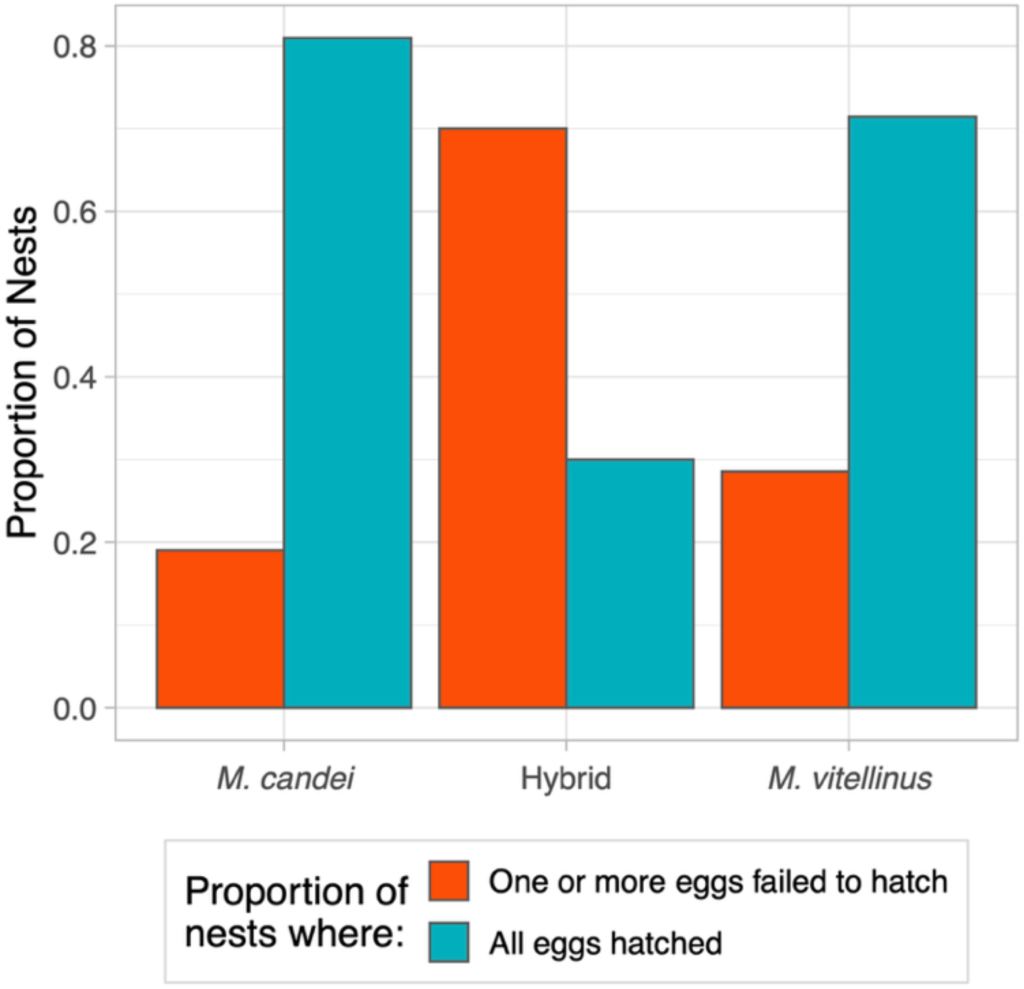
Paired bar plots of the frequency of nests in which: all eggs hatched successfully in a nest or one or both eggs failed to hatch within a nest across the three focal *Manacus* populations in the hybrid zone (N=21 *M. candei*), 10 (hybrid), and 7 (*M. vitellinus*) nests).

### Egg Candling Observations of Failed Eggs

At all three focal sites, 13 of the 16 eggs that failed to hatch never developed red vasculature (4 *M. candei*, 6 hybrid, 3 *M. vitellinus*) (Hatching Data: Table S3). Two of the failed eggs developed the characteristic red veining (1 *M. candei*, 1 hybrid) and this *M. candei* egg further developed a small embryo before ceasing development. For one egg from the hybrid population, we were unable to get candling observations during incubation because the female was on the nest during every 3-day nest check; however, when checking the nest after the 18-day incubation period, we found one nestling in the nest and the other egg was missing. We believe the female removed the egg after it failed to hatch. We observed this egg removal behavior twice in the hybrid focal population and once in *M. candei*. All three removals occurred in nests where two eggs were laid, and the failed egg disappeared after the successful egg hatched.

### Adult Survival

We captured 316 individuals from *M. candei* (N = 131), *M. vitellinus* (N = 65), and genomic center hybrids (N = 120), consisting of adult males (N = 111), breeding females (N = 102), and immature males, immature females, or non-breeding females of unknown age (N = 103) from February through June in the 2017, 2018, and 2019 sampling seasons. The apparent survival rates of hybrids (*φ* = 0.63, S.E. 0.099), and the parental species (*M. candei φ* = 0.44, S.E. 0.095, *M. vitellinus φ* = 0.50, S.E. 0.098) had extensive overlap in 95% confidence intervals (Figure 3); thus, we have little evidence that survival differs for hybrid individuals.

**Figure 3.**
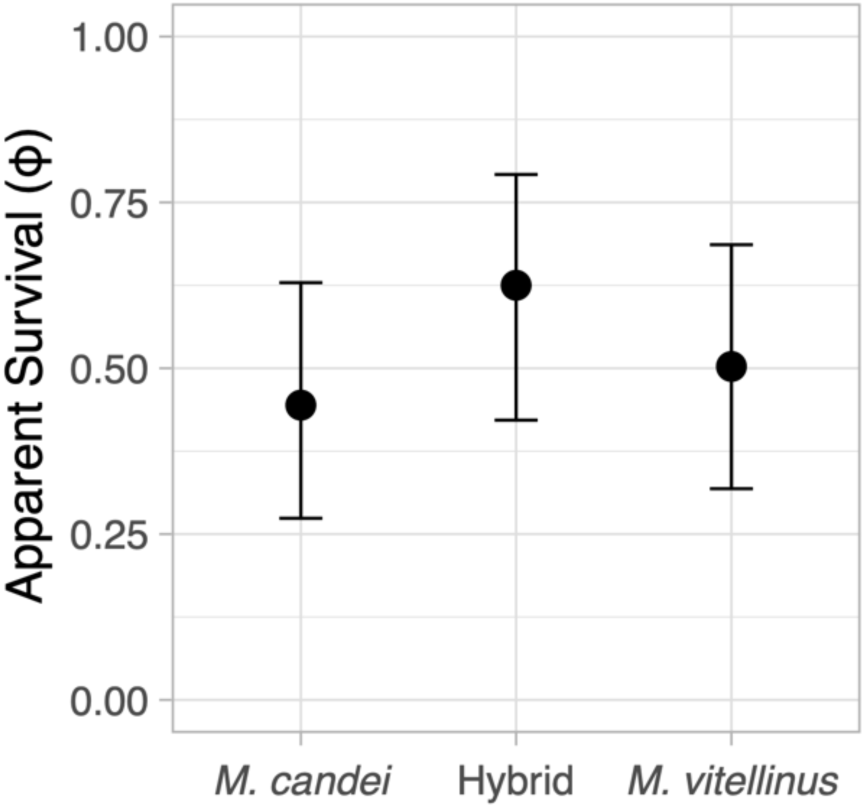
Estimated annual survival of hybrids and two parental species *Manacus* manakins with 95% confidence intervals. Estimates based on 316 individuals from *M. candei* (N = 131), *M. vitellinus* (N = 65), and hybrids (N = 120) from 2017-2019. We found no evidence of heterogeneity in recapture probability between the three groups: hybrids (p = 0.77, S.E. 0.13), *M. candei* (p = 0.69, S.E. 0.17) and *M. vitellinus* (p = 0.72, S.E. 0.16)

### Haemosporidian Parasitism

We screened 268 individuals for haemosporidian parasites (*Plasmodium* spp. and *Haemoproteus* spp.). Only 4 individuals tested positive for an active infection in all three PCR tests (Table 1). All infected individuals were *M. candei* (three adult males and one adult female) from the same sampling site (site 2, 8.33%, N = 48). Of all *M. candei* individuals tested (sites 2 and 3, N =73), 5.48% had active haemosporidian infections. No hybrid (N = 132) or *M. vitellinus* (N = 63) individuals tested positive for haemosporidian infection. Overall, from the 268 individuals assessed from across the hybrid zone, only 1.49% tested positive for a haemosporidian infection.

**Table 1.**
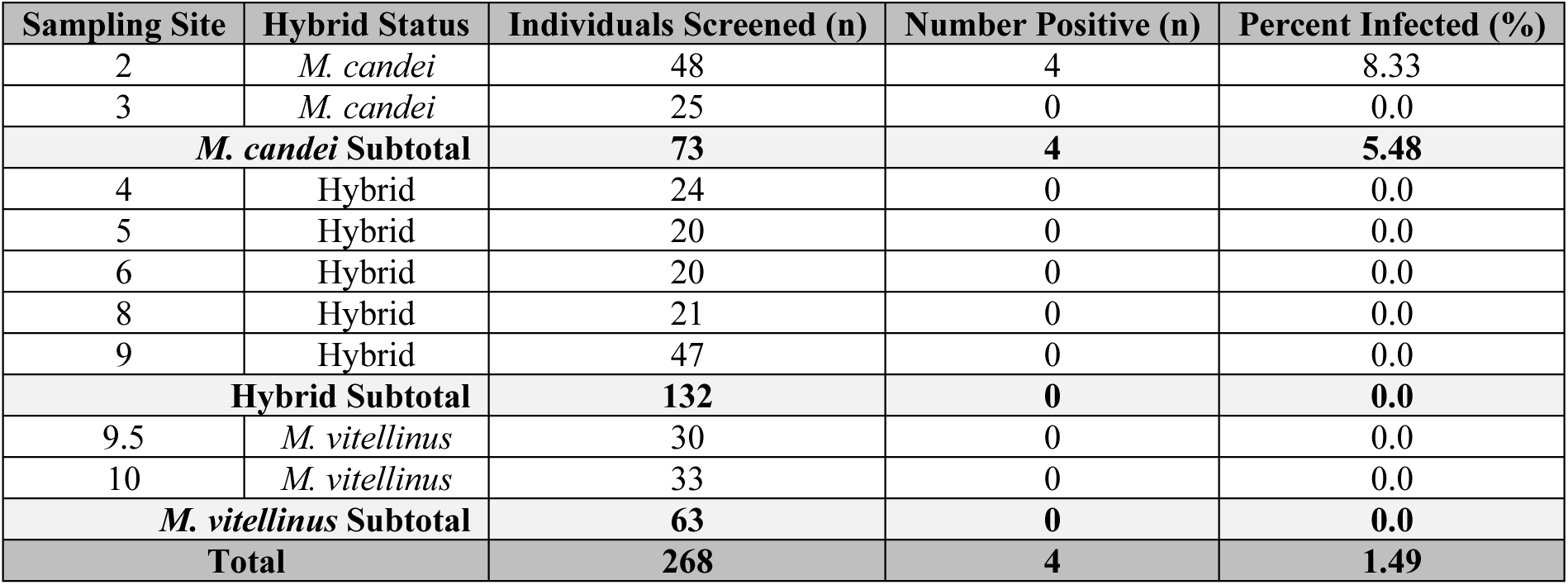
Sampling of *Manacus* individuals screened for haemosporidians and the number positive per sampling site. The four positive individuals were all *M. candei* and composed of 3 adult males and 1 adult female.

| Sampling Site | Hybrid Status | Individuals Screened (n) | Number Positive (n) | Percent Infected (%) |
| --- | --- | --- | --- | --- |
| 2 | *M. candei* | 48 | 4 | 8.33 |
| 3 | *M. candei* | 25 | 0 | 0.0 |
| ***M. candei* Subtotal** |  | **73** | **4** | **5.48** |
| 4 | Hybrid | 24 | 0 | 0.0 |
| 5 | Hybrid | 20 | 0 | 0.0 |
| 6 | Hybrid | 20 | 0 | 0.0 |
| 8 | Hybrid | 21 | 0 | 0.0 |
| 9 | Hybrid | 47 | 0 | 0.0 |
| **Hybrid Subtotal** |  | **132** | **0** | **0.0** |
| 9.5 | *M. vitellinus* | 30 | 0 | 0.0 |
| 10 | *M. vitellinus* | 33 | 0 | 0.0 |
| ***M. vitellinus* Subtotal** |  | **63** | **0** | **0.0** |
| **Total** |  | **268** | **4** | **1.49** |

## Discussion

Our findings show that reduced hatching success, but not reduced adult survival rates, is a consequence of hybridization in the *Manacus* hybrid zone. Nests from our hybrid focal population at the genomic center of the hybrid zone had significantly lower hatching success and a larger proportion of nests contained failed eggs when compared to the parental species.

Estimated adult survival rates were similar among focal populations. Additionally, we found the prevalence of infection by *Plasmodium* spp. and/or *Haemoproteus* spp. parasites was uniformly low across the entire hybrid zone. We posit that reduced reproductive success is the main avenue contributing to the many stable, narrow clines observed in the genomic center of the *Manacus* hybrid zone (Long et al. 2024).

### Reduced reproductive success signaling genetic incompatibilities

Reduced hatching success in nests from the hybrid population was the clearest evidence we found of a fitness cost to hybridization. Reduced hatching success (or nesting/fledging success as related proxies of reproductive success) has been observed in other hybrid zones and taxa (Kruuk et al. 1999, Svedin et al. 2008, Muñoz et al. 2010, Stelkens et al. 2015, Ålund et al. 2023), but is not universal (Vallender et al. 2007, Casas et al. 2012, Megna et al. 2014, Walsh et al. 2016). We found 13% (*M. candei*) and 23% (*M. vitellinus*) hatching failure, generally in line with the mean estimated rate of hatching failure in birds of 16.79% (Marshall et al. 2023). The proportion of hatching failure for nests in the hybrid focal population, however, was 42%, significantly higher than the parental species.

In birds, higher rates of hatching failure often appear in small populations where inbreeding is likely (Westemeier et al. 1998, Assersohn et al. 2021, Marshall et al. 2023). Marshall et al. 2023 found the highest rates of hatching failure in captive birds classified as threatened (by the IUCN Red List classifications) with 42.84% hatching failure. Threatened taxa are likely to be characterized by populations that are small, bottlenecked, or otherwise lacking genetic diversity. Inbreeding depression has been highlighted as one of the major drivers of hatching failure in threatened bird species (reviewed in Assersohn et al. 2021); however, we lack equivalent, systematic assessments of outbreeding depression/hybridization on hatching success. Inbreeding depression is generally due to decreasing heterozygosity, thereby increasing the likelihood of deleterious alleles coming to fixation at homozygous sites, conversely, outbreeding depression is more likely due to breaking associations of coadapted alleles at multiple loci across the genome (Shields 1982). While inbreeding and outbreeding depression are seemingly opposites, the shared phenotype of hatching failure invites inquiry into whether inbreeding and outbreeding are tapping into similar developmental processes and if additional studies of hybrid hatching failure could help elucidate common drivers of hatching failure.

The stage of development at which an embryo dies can offer clues as to which biological processes are going awry and guide further investigations. Embryo mortality in birds, in general, occurs in either the early or late stages of development before hatching (Romanoff 1949, Assersohn et al. 2021). Early embryo mortality (e.g., less than 72 hours after successful fertilization), in particular, is commonly associated with genetic problems (Shook et al. 1971, Assersohn et al. 2021). Experimental work in inter-genic hybrids between chickens (*Gallus gallus domesticus*) and Japanese quail (*Coturnix japonica*), which have extremely high hatching failure in F1 hybrids and all resulting hybrid males are sterile, found that 75-80% of hybrid embryos ceased development by the second day of incubation (Ishishita et al. 2016). Failure in these first two incubation days means that embryos stopped developing before the formation of discernible embryonic structures, including extraembryonic membrane and blood island formation (Ishishita et al. 2016). Thus, these sorts of common early embryonic egg failures may appear like unfertilized eggs when using the field candling techniques we employed.

While we were unable to predictably monitor *Manacus* eggs from their first few hours of incubation, all hybrid eggs that failed to hatch by the end of the incubation period did not form an embryo large enough to detect with our field candling (they all retained a clearly defined yellow yolk or had bright red vascularization when illuminated), suggesting that either fertilization failure occurred or embryo development was arrested in the early stages when the embryo was very small (i.e., within the first two days). However, all but a single *M. candei* egg from the parental species also failed to develop a visible embryo, although there were fewer failed eggs to observe in the parental species. Nevertheless, without using microscopy to examine the eggs that failed to hatch, we cannot determine if these eggs failed due to fertilization failure or early embryonic death, which would each point to different underlying mechanisms.

For example, if hybrid eggs truly have higher fertilization failure, this could imply the underlying mechanism is related to deficiencies in hybrid sperm such as low quantities or poor sperm function. However, if hybrid eggs were failing from early embryonic death in the first few days of incubation, this would point to lethal problems in developmental processes such as protein synthesis, cell proliferation, and gastrulation, which all occur in intergenic chicken-quail hybrids from misregulated gene expression in these same pathways (Ishishita et al. 2020). Future studies should candle eggs to monitor egg development, collect failed eggs to determine the exact cause of hatching failure, and/or collect all eggs and utilize artificial incubation to detail embryo development in a controlled environment to determine exactly when and how hybrid embryos cease development. Artificial incubation has been successfully done in manakins (Jones and DuVal 2019), and could be an important tool for future investigations of hybrid hatching success.

Genetic incompatibilities between the two parental species, e.g., Bateson–Dobzhansky– Muller incompatibilities (Bateson 1909, Dobzhansky 1936, Coyne and Orr 2004), are a likely mechanism underlying the reduced hatching success in the *Manacus* hybrid zone. The genetic center of the hybrid zone may be acting as a filter (Martinsen et al. 2001) taking out incompatible alleles early on in development and leaving only hybrid adults with the successful allelic combinations. We observed similar rates of adult apparent survival between the hybrid and parental focal groups, consistent with this idea. We find further evidence from Vernasco et al. (2024), who observed that birds sampled at site 9 have longer telomeres and increased heterozygosity in comparison to parental populations at sites 2 and 10. While the specific fitness effects of telomere lengths remain unknown, longer telomere lengths are often associated with higher fitness, providing preliminary support for hybrid manakins having equal-to-higher fitness than the parents once individuals with incompatible alleles are removed during development.

Other avenues of selection should be explored as well to assess additional selective pressures on hybrid populations, such as metabolic or physiological robustness in hybrid individuals that could affect their courtship displays or dispersal ability. Moreover, collecting failed hybrid eggs is ripe for genetic studies to uncover unsuccessful allelic combinations seen only in unhatched hybrids, and these genes are likely vitally important for the speciation process (Presgraves 2010).

Reduced hatching success could be a particularly strong selective pressure for hybrid zone dynamics in tropical species, given generally small clutch sizes (1-2 eggs) and high rates of nesting failure (Brawn et al. 2011, Stutchbury and Morton 2023). If, for example, a temperate bird lays a 5-8 egg clutch and half fail to hatch from hybrid infertility or inviability, then that nest could still produce 3-4 successful offspring (as seen in Bronson et al. 2005). In contrast, many tropical birds, including manakins, lay only 2 eggs, which would result in only one successful offspring if half fail to hatch. While tropical species tend to renest more often due to longer breeding seasons (Stutchbury and Morton 2023), higher hatching failure from genetic incompatibilities would likely still be present in subsequent renesting attempts. Thus, hybrid hatching success could be a key factor in regulating avian hybrid zone dynamics in the tropics, and more studies on tropical hybrid zones could confirm if this is a common mechanism across taxa and latitudes.

### Haemosporidia as a selective pressure in hybrid zones and manakins

Parasitism has been proposed as an important mechanism for shaping hybrid zones (reviewed in Theodosopoulos et al. 2019); however, there are mixed results on the effects of avian blood parasites on hybridization dynamics (Reullier et al. 2008, Cozzarolo et al. 2018, Roth et al. 2021, Rice et al. 2021). We found only 1.49% positive infections for haemosporidian parasites across the entire *Manacus* hybrid zone in western Panama, indicating that *Plasmodium* and *Haemoproteus* do not play a major role in the hybridization dynamics of our focal system. This result is somewhat surprising, given the ubiquitous prevalence of haemosporidian parasites in other neotropical species such as clay-colored thrush (*Turdus grayi*) (Ricklefs and Sheldon 2007) and bananaquit (*Coereba flaveola*) (Ricklefs et al. 2011, Antonides et al. 2019), both of which are commonly found adjacent to *Manacus* leks.

In other manakin species, the percent prevalence of haemosporidian infections such as avian malaria tends to be low compared to the rest of the avian community (when surveyed), but there are mixed results depending on the species assessed. For example, blue-crowned manakins (*Lepidothrix coronata*) had 35% prevalence (Bosholn et al. 2016), while the wire-tailed manakins (*Pipra filicauda*) had only 5.6%, the lowest prevalence in an assemblage of 39 species (Svensson-Coelho et al. 2013). Across 30 species of manakins only 22 had any malaria detected, with an average prevalence of only 10.9% (ranging from 0-50%). Yet, *Manacus* maintained some of the lowest malaria prevalence with *Manacus vitellinus* at 7.5% and *Manacus manacus* at only 6.7% (Fecchio et al. 2017). Previous studies on *Manacus* manakins have all found low levels of haemosporidian infection, including zero infections detected in *M. candei* (Valkiūnas et al. 2004) and 2 out of 6 infected *M. vitellinus* individuals (Ricklefs and Sheldon 2007), but sample sizes to date have been quite small. Our larger sampling from *M. candei* and *M. vitellinus* does, however, corroborate these previous findings that *Manacus* have low haemosporidian prevalence.

We hypothesize that manakins, especially *Manacus*, might not readily contract haemosporidian infections. However, we did not collect blood samples from non-target species on the leks, so we cannot say for certain if *Manacus* manakins specifically have abnormally low prevalence rates compared to the local avian communities around secondary forests in Bocas del Toro, Panama. As far as we know, prevalence of avian haemosporidian parasites has not been previously estimated in the province of Bocas del Toro; future studies of blood parasite infections at the community level will clarify whether low prevalence is systemic among species in the region’s bird communities, or specific to *Manacus* manakins.

## Conclusion

We explored hybrid viability in a tropical, avian hybrid zone governed by strong selection on male secondary sexual traits. We assessed putative hybrid fitness by investigating adult apparent survival and hatching success along with a prospective selective pressure of parasitism by haemosporidian parasites. We found that hybrids have reduced hatching success, but no evidence of reduced survival or susceptibility to avian malarial infections. Thus, we believe that hybrid dysfunction is manifesting in this hybrid zone through reduced reproductive success, and this could be contributing to hybrid zone stability. Linking reduced hatching success to specific genetic incompatibilities in hybrid genomes is an important next step to uncovering the dynamics of hybridization and speciation.

## Acknowledgments

We thank the Smithsonian Tropical Research Institute and associates including Owen McMillan, Adriana Bilgray, Lil Camacho, Isis Ochoa, Juan Maté and MiAmbiente for permits and logistics. We also thank Cesar Romero for extensive logistical support. Our field assistants who provided long hours of hard work: André Nguyen, Alexander Worm, Ivy Ciaburri, Olivia Ferrari, Elena Prado-Ragan, Vaughn Hage, Luis Ramos Vazquez, and Evalynn Trumbo, as did our lab assistant Bingting (Grace) He. We would also like to thank the families and local communities who generously permitted us to work on their land including but not limited to the Jimenez Family, the Lezcano family, the Aguilar Family, the López family, and the Naso and Ngäbe-Buglé peoples of Bocas del Toro. Thank you to the staff at the University of Illinois at Urbana-Champaign departments of Natural Resources and Environmental Sciences and Integrative Biology, along with Judith Toms, Julian Catchen, Becky Fuller, Angel G. Rivera-Colón, Todd Jones, Emma Young, Karthik Yarlagadda, and George Pantazes, for helpful discussions on the manuscript, analyses, and protocols. This project was supported by the University of Illinois Urbana-Champaign, the NSF DGE 10-69157 IGERT to KML, the USDA National Institute of Food and Agriculture Hatch project 1026333 (ILLU-875-984), and the Manakin Research Coordination Network (NSF DEB 1457541).

## Data Availability Statement

Scripts required for running the apparent survival and egg hatching statistics are available in the following repository: <u>github.com/kiralong/manacus_HZ_fitness_ms</u>. Raw nesting, hatching, and malaria data will be provided as a supplementary excel file with the final publication (“Supplementary_Data.xlxs: S1.Raw_capture_record, S2. Molecular_sexing_results, S3.Hatching_data, and S4.Malaria_data”).

## Appendix

**Table A1:**
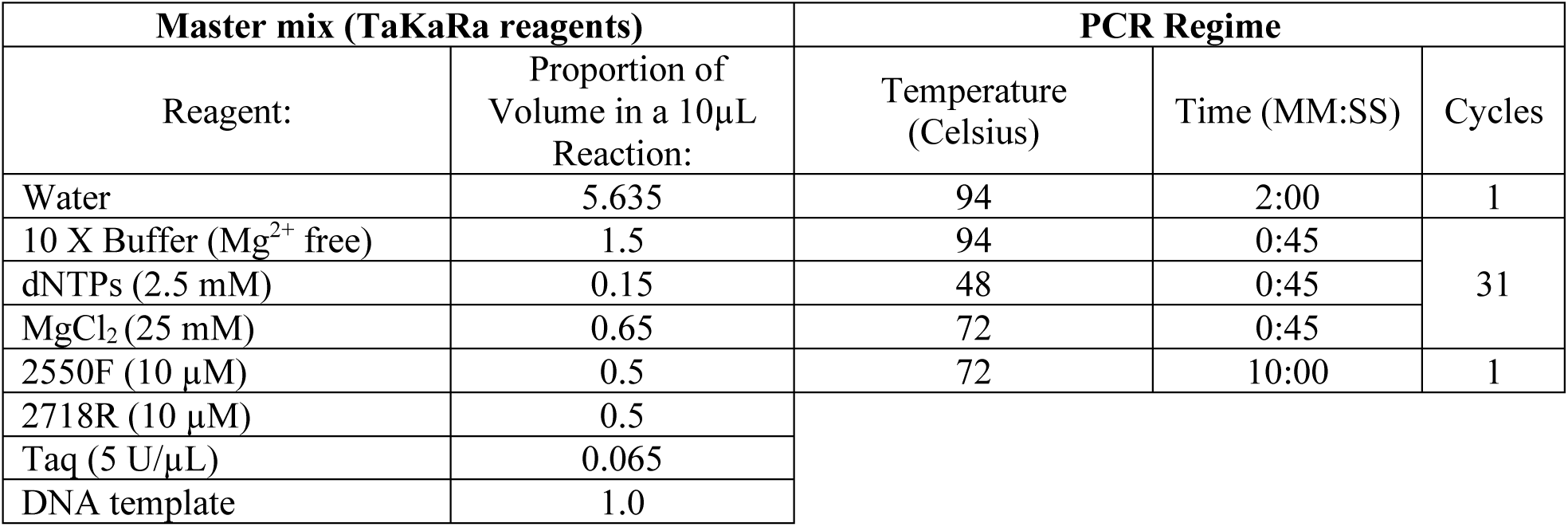
Molecular Sexing Protocol. Specific methods for the master mix and PCR cycles to identify molecular sex in birds. TaKaRa reagents are the standard kit concentrations from TaKaRa Ex Taq® (Mg2+ free Buffer) kit (Catalog # RR01AM). The PCR protocol and primers 2550F and 2718R are based on Fridolfsson and Ellegren (1999).

| Master mix (TaKaRa reagents) |  | PCR Regime |  |  |
| --- | --- | --- | --- | --- |
| Reagent: | Proportion of Volume in a 10µL Reaction: | Temperature (Celsius) | Time (MM:SS) | Cycles |
| Water | 5.635 | 94 | 2:00 | 1 |
| 10 X Buffer (Mg<sup>2+</sup> free) | 1.5 | 94 | 0:45 | 31 |
| dNTPs (2.5 mM) | 0.15 | 48 | 0:45 |  |
| MgCl<sub>2</sub> (25 mM) | 0.65 | 72 | 0:45 |  |
| 2550F (10 µM) | 0.5 | 72 | 10:00 | 1 |
| 2718R (10 µM) | 0.5 |  |  |  |
| Taq (5 U/µL) | 0.065 |  |  |  |
| DNA template | 1.0 |  |  |  |

**Table A2:**
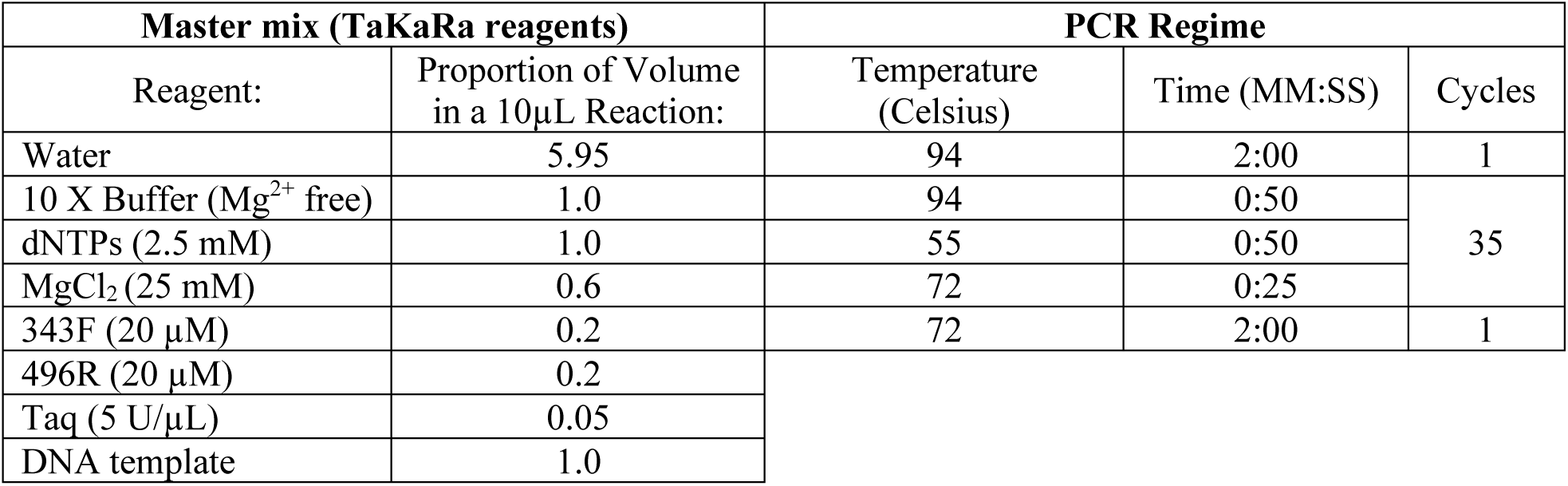
Molecular malaria screening protocol for master mix and PCR cycles. TaKaRa reagents are the standard kit concentrations from TaKaRa Ex Taq® (Mg2+ free Buffer) kit (Catalog # RR01AM). The PCR protocol and primers *343F* and *496R* are based on Fallon et al. (2003).

**Table A3:**
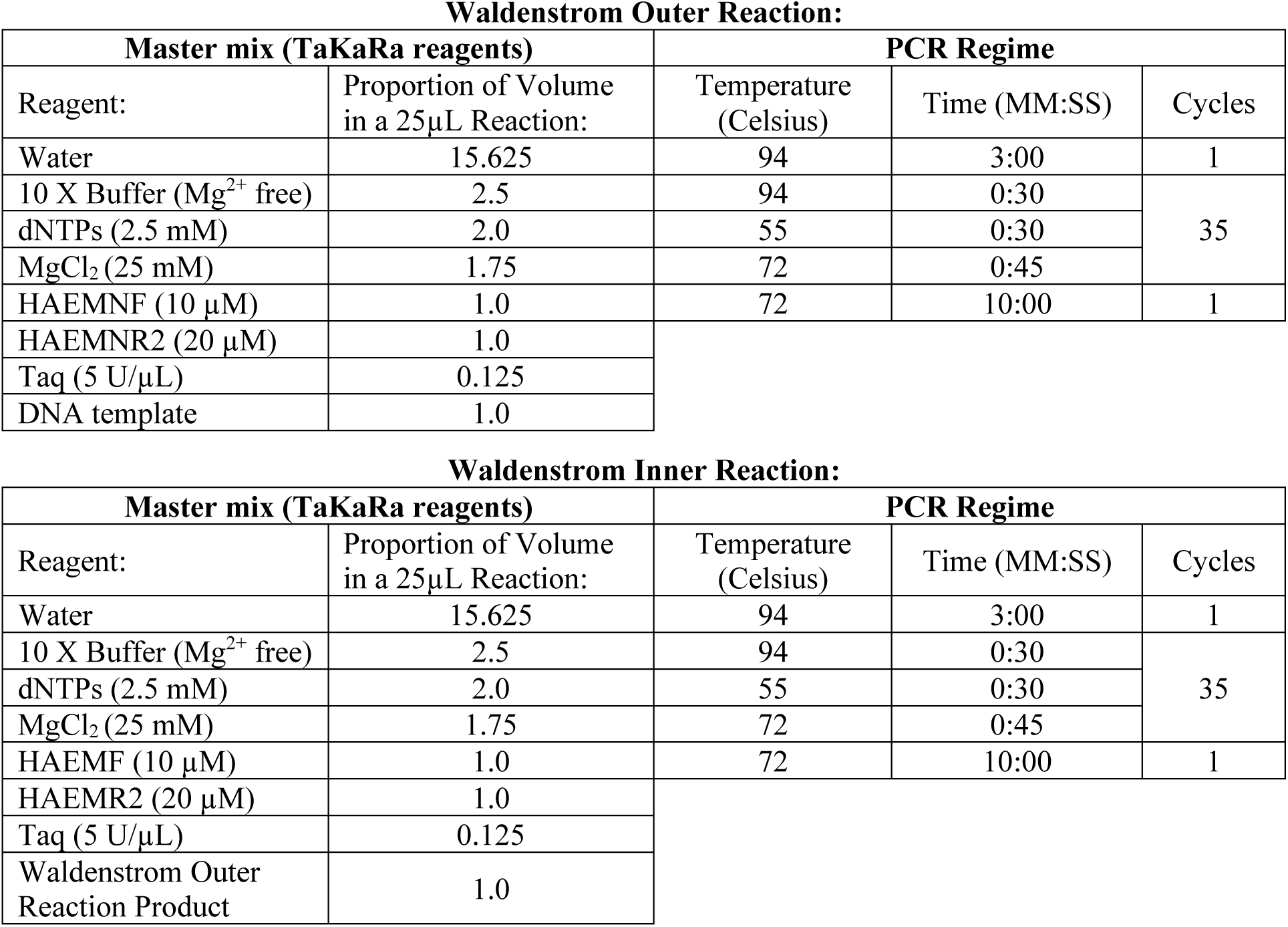
Molecular Malaria Waldenstrom Protocol, a nested PCR protocol to detect haemosporidians. TaKaRa reagents are the standard kit concentrations from TaKaRa Ex Taq® (Mg2+ free Buffer) kit (Catalog # RR01AM). For this nested PCR assay, we used the *HAEM* primers from Waldenstrom et al. (2002).

**Waldenstrom Outer Reaction:**
| Master mix (TaKaRa reagents) |  | PCR Regime |  |  |
| --- | --- | --- | --- | --- |
| Reagent: | Proportion of Volume in a 25µL Reaction: | Temperature (Celsius) | Time (MM:SS) | Cycles |
| Water | 15.625 | 94 | 3:00 | 1 |
| 10 X Buffer (Mg<sup>2+</sup> free) | 2.5 | 94 | 0:30 | 35 |
| dNTPs (2.5 mM) | 2.0 | 55 | 0:30 |  |
| MgCl<sub>2</sub> (25 mM) | 1.75 | 72 | 0:45 |  |
| HAEMNF (10 µM) | 1.0 | 72 | 10:00 | 1 |
| HAEMNR2 (20 µM) | 1.0 |  |  |  |
| Taq (5 U/µL) | 0.125 |  |  |  |
| DNA template | 1.0 |  |  |  |

**Waldenstrom Inner Reaction:**
| Master mix (TaKaRa reagents) |  | PCR Regime |  |  |
| --- | --- | --- | --- | --- |
| Reagent: | Proportion of Volume in a 25µL Reaction: | Temperature (Celsius) | Time (MM:SS) | Cycles |
| Water | 15.625 | 94 | 3:00 | 1 |
| 10 X Buffer (Mg<sup>2+</sup> free) | 2.5 | 94 | 0:30 | 35 |
| dNTPs (2.5 mM) | 2.0 | 55 | 0:30 |  |
| MgCl<sub>2</sub> (25 mM) | 1.75 | 72 | 0:45 |  |
| HAEMF (10 µM) | 1.0 | 72 | 10:00 | 1 |
| HAEMR2 (20 µM) | 1.0 |  |  |  |
| Taq (5 U/µL) | 0.125 |  |  |  |
| Waldenstrom Outer Reaction Product | 1.0 |  |  |  |

## Notes

https://github.com/kiralong/manacus_HZ_fitness_ms

### Competing Interest Statement

The authors have declared no competing interest.

